# Nuclear localization of calcium-activated BK channels in skate ampullary electroreceptors

**DOI:** 10.1101/2020.01.27.922161

**Authors:** Abby L. Chen, Ting-Hsuan Wu, Lingfang Shi, William T. Clusin, Peter N. Kao

## Abstract

Ampullae of Lorenzini are sensory organs capable of detecting microvolt gradients in seawater. Electroreception involves interplay between voltage-dependent calcium channels Ca_V_1.3 and big conductance calcium-activated potassium (BK) channels in apical membranes of receptor cells. Expression of BK (*kcnma1*) and Ca_V_1.3 (*cacna1d*) channels in skate (*Leucoraja erinacea)* ampullary electroreceptors was studied by *in situ* confocal microscopy. BK and Ca_V_1.3 channels colocalize in plasma membranes, ribbon synapses and kinocilia. BK channels additionally colocalize with chromatin and nuclear lamins in electroreceptor cells. Bioinformatic sequence analysis identified an alternatively spliced bipartite nuclear localization sequence (NLS) in *kcnma1* (at site of mammalian STREX exon). Skate *kcnma1* wild type cDNA transfected into HEK293 cells localized to the endoplasmic reticulum and nucleus. Mutations in the NLS (KR→AA or SVLS→AVLA) independently attenuated nuclear translocation from endoplasmic reticulum. BK channel localization may be controlled by splicing or phosphorylation to tune electroreception and modulate gene expression.

## Introduction

Calcium-activated potassium channels were first discovered in the early 1970’s and are present in virtually all living organisms. The physiological roles ascribed to these channels have greatly expanded since the well-known review article by Meech in 1978 [1]. Dysfunction of calcium-activated potassium channels are involved in a broad range of neurologic, cardiac and autoimmune diseases, including epilepsy [2], alcoholism [3], cardiac arrhythmias [4, 5], rheumatoid arthritis [6–8] and skeletal myoblast differentiation [9].

Ampullae of Lorenzini are electroreceptive sensory organs capable of detecting nanovolt gradients in seawater [10, 11]. Canals filled with highly electroconductive keratan sulfate jelly [12] originate anterior to gill slits and terminate in ampullary outpouchings of epithelia, from which the sensory nerve derives. Each ampulla opens into seven alveoli consisting of epithelial sheets of conical electroreceptor cells with apical kinocilia facing the lumen, intercalated between supporting cells that form electrically tight junctions to the receptor cells [13, 14].

The types and location of ion channels in the receptor cells was inferred from anatomical, physiological and pharmacologic data [15–20]. In response to an excitatory electrical stimulus (when the lumen becomes more positive), opening of voltage-dependent calcium channels produces a depolarizing inward calcium current entering the receptor cell through the apical membrane, that flows outward across the basal membranes, triggering calcium-dependent vesicle release. The excitatory response is terminated by opening of calcium-activated potassium channels in the apical membrane, which repolarizes both faces [21–25]. Spontaneous activity is characterized by 20 Hz oscillations that can be recorded experimentally by placing the ampulla in an air gap [18, 22]. Voltage-gated potassium channels in the receptor cells contribute to the oscillatory responses [19, 20]. These oscillatory signals control release of neurotransmitter inferred to be glutamic acid by RNA-ISH localization of Slc17a8 (glutamate transporter 3) [26].

Release of calcium from intracellular stores can also lead to opening of calcium-activated potassium channels. There are two principal types of calcium-activated potassium channels. Small conductance calcium activated potassium channels (SK) have a single channel conductance of 9-14 pS and large conductance channels (BK), have a conductance of 250-300 pS [27, 28]. BK channels are generally sensitive to both intracellular calcium and membrane voltage, while SK channels are only sensitive to calcium.

Recent studies by Bellono *et al.* determined the amino acid sequences for Ca_V_1.3 *cacna1d* and BK *kcnma1* channels in skate electroreceptors [29], suggesting that a slight sequence variation in the calcium channel in the skate makes it unusually sensitive to voltage. Relative expression of mRNA transcripts for BK channels was 35-fold higher than for SK channels. Furthermore, *in situ* hybridizations demonstrated high expression of mRNA transcripts for *cacna1d* and *kcnma1* in electroreceptor cells of skate ampulla.

Trafficking of potassium channels to intracellular organelles or plasma membranes is subject to complex regulation that includes alternative splicing to incorporate specific addresses, posttranslational modifications such as phosphorylation or glycosylation and association with accessory subunits [30, 31]. The voltage-gated K channel, Kv10.1 is linked to diverse cancers [32]. Functional Kv10.1 channels localized to the inner nuclear membrane together with Lamin A/C subsequent to exposure to the extracellular milieu [33].

BK channels are regulated by complex alternative splicing, post-translational modifications and protein-protein interactions that together influence subcellular localization and function [27, 34–44]. Alternative splicing at the C-terminus that incorporates amino acids -DEC targets BK channels to mitochondria [45]. Complex splicing at site 4, also known as c2, where the mammalian STREX exon may insert, was shown to affect BK*α* subunit channel tissue distribution, trafficking and regulation [36, 38]. Random insertion of GFP variants into BK channels resulted in generation of 55 unique fluorescent fusion proteins, of which 19 were expressed and functional at the plasma membrane and 36 showed intracellular expression without suggestion of misfolding [46].

Nuclear localization of calcium-activated BK channels has been described [47–50]. Li et al [51] demonstrated BK channel expression in hippocampal neurons in a pattern of intracellular rings surrounding nuclei and colocalizing with the nuclear envelope marker lamin B1. Confocal microscopy was used to demonstrate colocalization of BK channels with lamin B1 in inner nuclear membranes. These signals were absent in neurons from mice with targeted knockout of BK*α*.

Skate ampullae are small and translucent making them suitable for whole mount immunostaining and *in situ* confocal microscopy. We studied skate electroreceptor expression of voltage-gated calcium channel Ca_V_1.3 and calcium-activated potassium channel BK using antibodies generated against conserved epitopes. Unexpectedly, we discovered prominent nuclear expression of BK channels in electroreceptor cells.

## Results

Skate ampullary electroreceptors are thin walled epithelial structures that transduce microvolt electrical gradients into membrane oscillations that regulate synaptic transmission. Ampulla with attached canals and sensory nerves were dissected from little skates, *Leucoraja erinacea*. Freshly dissected ampulla were fixed in paraformaldehyde, permeabilized, and used for whole mount immunostaining. Primary antibodies were selected based on sequence conservation of epitopes between skate and mammalian BK and Ca_V_1.3 channels. Nuclei were counterstained with DAPI. Confocal z-stacks of 1-2 micron optical sections spanning up to 100 microns were acquired at 40x magnification through individual alveoli of the ampulla.

A high magnification optical section across the involuted wall of an alveolus is selected to illustrate notable features of BK channel *α* subunit expression and localization (Figure 1; Supplemental Movie 1). Anti-BK*α* rabbit polyclonal antibody PA1-923 shows strong reactivity in disks (Figure 1A) that colocalize with DAPI nuclear staining of chromatin (Figure 1B, C). The nuclear expression of BK detected by PA1-923 varies across the alveolar wall: the strongest BK expression in nuclear disks is present in cells one layer removed from the alveolar lumen (Figure 1C, gradient of pink). BK expression was also detected in hexagons, which we interpret as plasma membrane expression captured in cross-section (Figure 1A, C). We observed that cells with prominent hexagonal plasma membrane BK expression exhibited relatively lower levels of nuclear BK expression (Figure 1A). We independently detected BK expression using mouse monoclonal antibody MaxiK*α* (Figure 1D). MaxiK*α* also detected BK expression in disks (colocalizing with DAPI) and showed stronger immunoreactivity than PA1-923 in punctate foci and short spikes pointing into the alveolar lumen, which we interpret to represent BK channel expression in sensory kinocilia (Figure 1D-F, “k”). MaxiK*α* also showed stronger reactivity than PA1-923 at the basal, outer surfaces of alveoli where ribbon synapses of electroreceptor cells connect to the afferent nerves (Figure 1D-F, “rs”).

**Figure 1:**
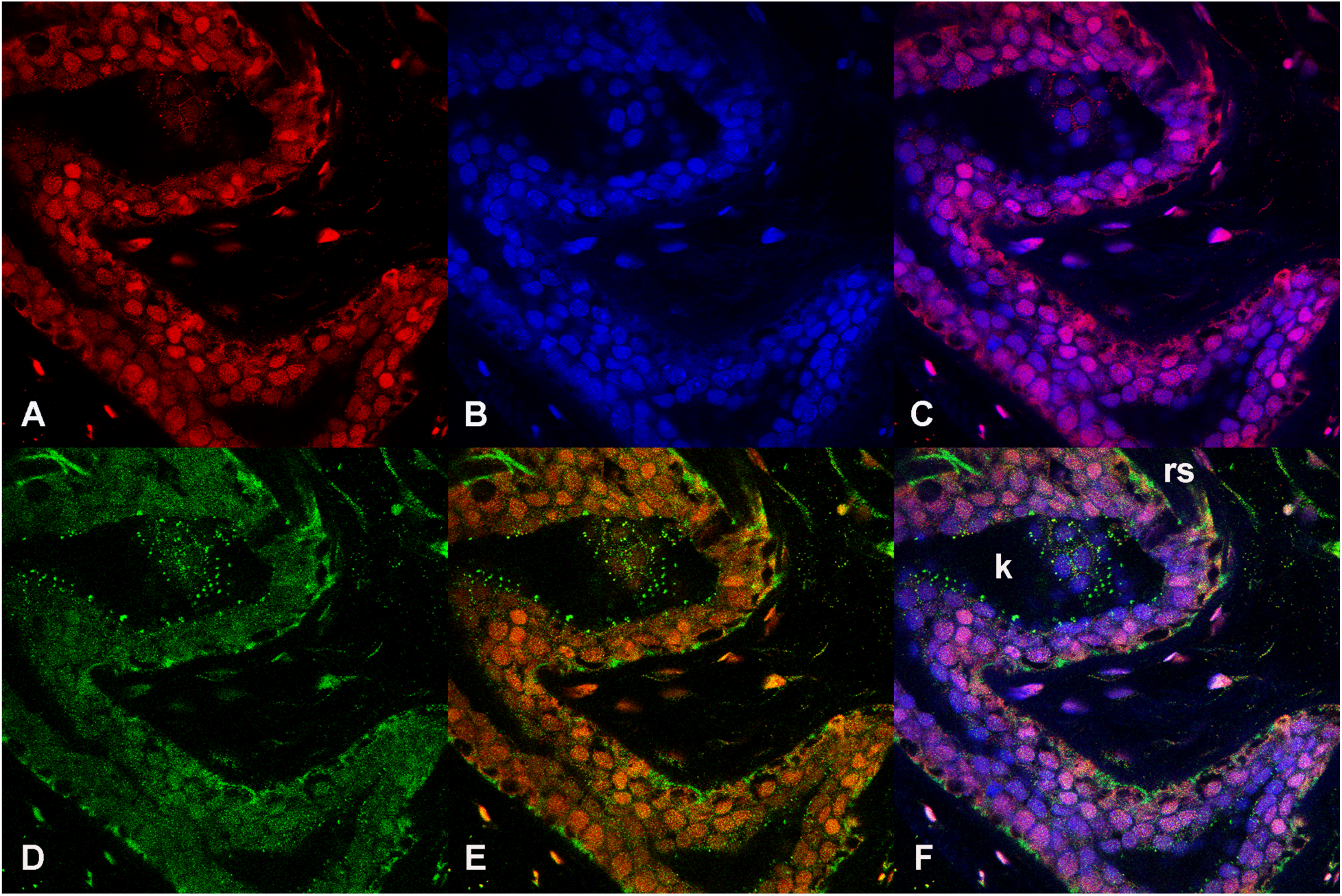
BK channels are present in nuclei and plasma membranes of skate ampullary electroreceptor. A) BK antibody PA1-923 (rabbit polyclonal). B) DAPI nuclear stain. C) Merge of A and B. D) BK antibody MaxiK*α* (mouse monoclonal). E) Merge of A and D. F) Merge of A, B and D.

Colocalization of BK channels with nuclear lamins was next investigated. We combined rabbit anti-BK (PA1-923) with mouse anti-lamin A/C (Figure 2; Supplemental Movie 2). BK immunoreactivity was prominent in nuclear disks (Figure 2A,C,F). Lamin A/C immunoreactivity was present in disks (Figure 2D) that colocalized with DAPI nuclear staining (Figure 2B) and BK expression (Figure 2E,F). We detected lamin B1 using a rabbit antibody and combined this with anti-BK mouse MaxiK*α* (Figure 3; Supplemental Movie 3). BK immunoreactivity was prominent in rings (Figure 3A) surrounding nuclei (Figure 3B,C,F). Lamin B1 immunoreactivity marks inner nuclear membranes as rings (Figure 3D) and colocalizes precisely with BK immunoreactivity (Figure 3E,F) as showed by the pixel color shift of green + red to yellow. The arrow (Figure 3F) marks one of several puncta with strong BK and moderate lamin B1 immunoreactivity, which we interpret to represent the base of a kinocilia where it merges with the nuclear envelope (Supplemental Movie 3).

**Figure 2.**
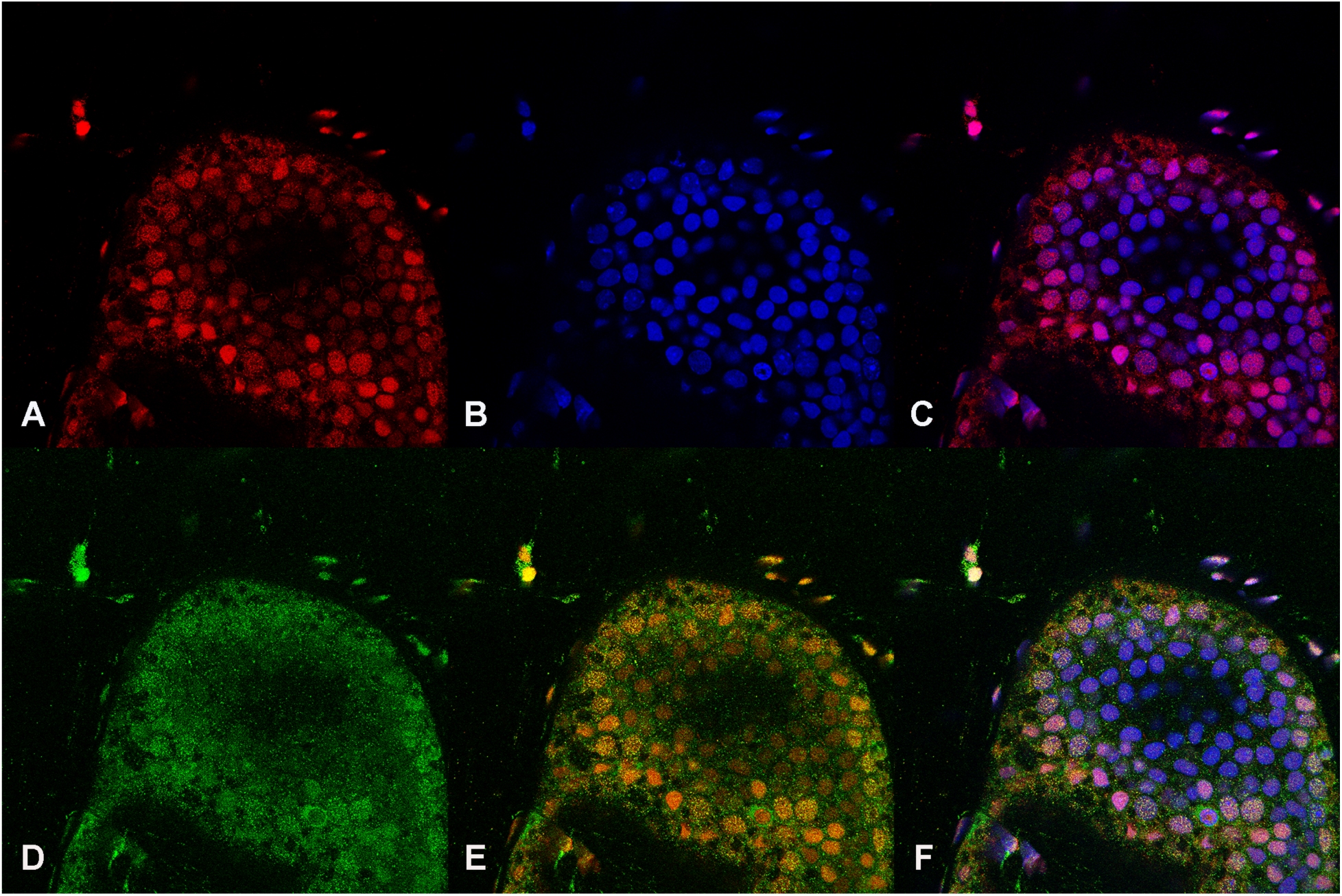
BK channels colocalize with Lamin A/C. A) BK antibody (PA1-923, rabbit). B) DAPI. C) Merge of A and B. D) Lamin A/C antibody (mouse monoclonal). E) Merge of A and D. F) Merge of A, B and D.

**Figure 3.**
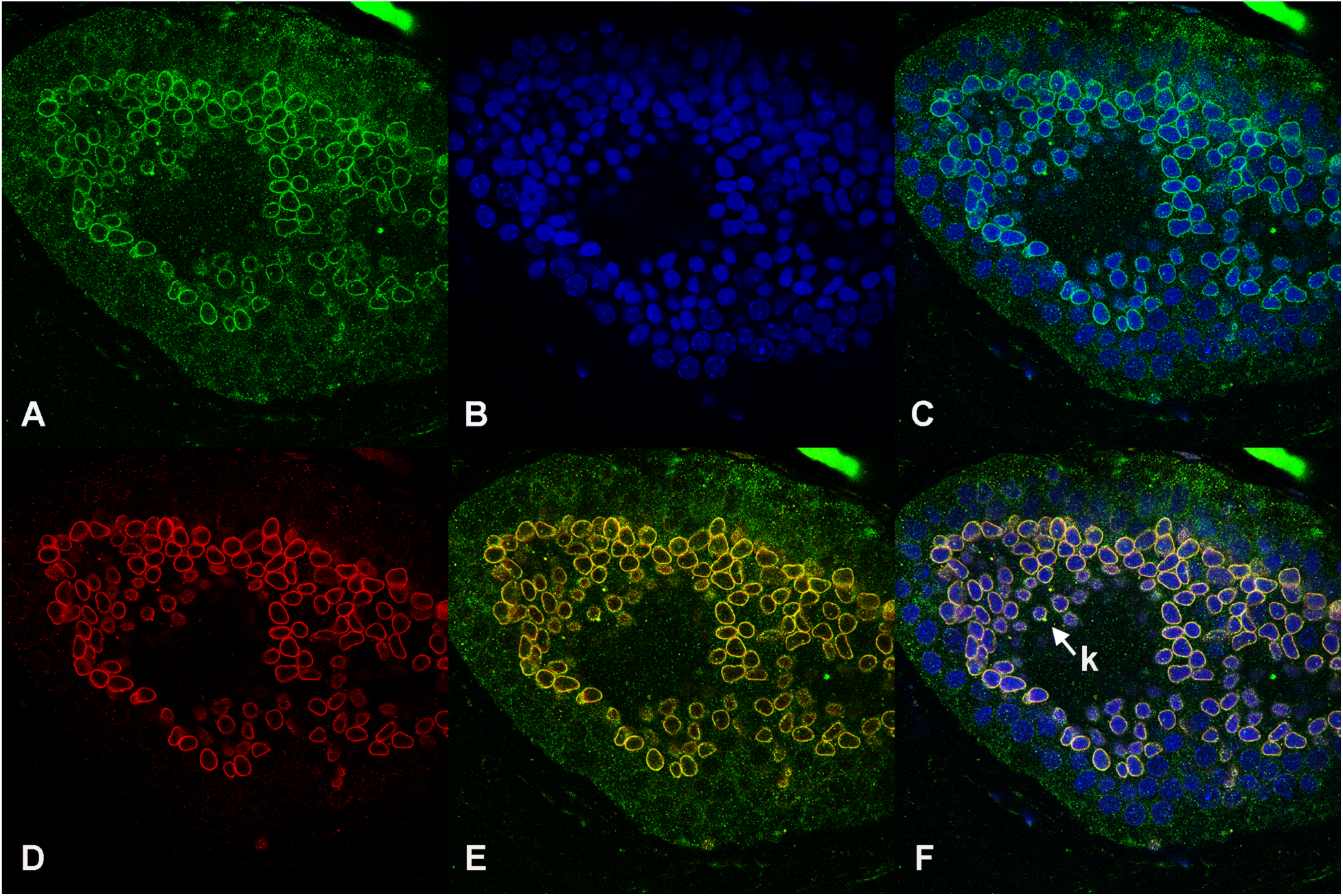
BK channels colocalize with Lamin B1. A) BK antibody (MaxiK*α*, mouse monoclonal). B) DAPI. C) Merge of A and B. D) Lamin B1 antibody (rabbit polyclonal). E) Merge of A and D. F) Merge of A, B and D. “k” marks putative kinocilia base.

Expression of Ca_V_1.3 was determined using mouse antibody LS-B4915 raised against rat Ca_V_1.3 amino acids 859-875 (Figure 4; Supplemental Movies 5, 6). Skate Ca_V_1.3 has 11/17 conserved amino acids in this epitope. Ampullae were coimmunostained with mouse anti-Ca_V_1.3 (Figure 4A,G) and rabbit anti-BK*α* (PA1-923) (Figure 4D,J) and nuclei were stained with DAPI (Figure 4B,H). Ca_V_1.3 is expressed in plasma membranes, ribbon synapses and kinocilia (Figure 4A,F, G,K; “rs”, “k”). In contrast to BK channels (Figure 4D,J), there is little nuclear expression of Ca_V_1.3. Colocalization of Ca_V_1.3 and BK channels is seen in plasma membranes (Figure 4E,K). Ca_V_1.3 expression is higher than BK expression at the outer, basal surfaces of electroreceptor cells where the ribbon synapses give rise to the afferent nerve fibers (Figure 4F,L; “rs”). Colocalization of Ca_V_1.3 and BK channels is also concentrated at distinct punctae or short spikes which we interpret as kinocilia (Figure 4F,L, “k”). BK expression in disks overlying nuclei is seen in both the “wall” (Figure 4D-F) and “dome” (Figure 4J-L) views. In this specimen showing the alveolar “wall” (Figure 4A-F), there is again evidence of increasing BK nuclear expression in the layer of electroreceptor cells farthest from the lumen (Figure 4D-F). Additionally, the DAPI staining pattern differs between the two layers of nuclei in the wall of the alveolus (Figure 4B): those cells with greatest BK nuclear expression farthest from the lumen show mottled DAPI staining intercalated with BK staining (Figure 4F), whereas the nuclei closest to the lumen show uniform fine granular DAPI staining and little BK expression (except for single red dots). We propose the nuclear BK expression near chromatin may exist in nucleoplasmic envelopes connected to a nucleoplasmic reticulum [52], and contribute to regulation of gene expression, as previously reported [51]. Ultrastructural studies of Ampulla of Lorenzini by electron microscopy revealed that electroreceptor kinocilia intercalate between supporting cells that face the alveolar lumen [14].

**Figure 4:**
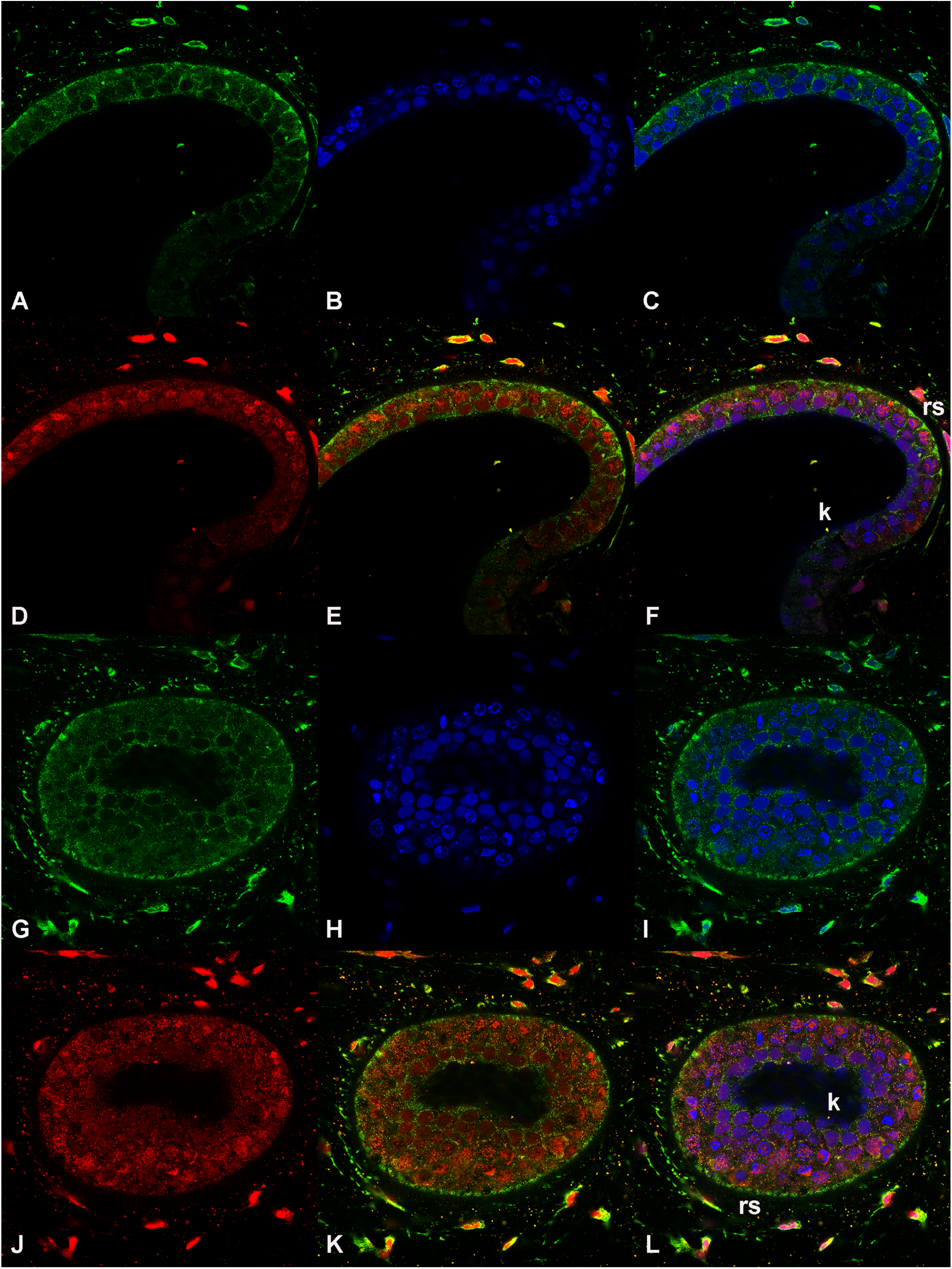
Expression of Ca_V_1.3 and BK channels in skate ampullary electroreceptor. Wall A-F, Dome G-L. A,G) Ca_V_1.3 antibody (mouse monoclonal LS-B4915). B,H) DAPI. D,J) BK antibody (rabbit polyclonal PA1-923). C) Merge of A and B. E) Merge of A and D. F) Merge of A, B and D. I) Merge of G and H. K) Merge of G and J. L) Merge of G, H and J. “k” adjacent to luminal kinocilia, “rs” adjacent to basal ribbon synapses.

We analyzed skate BK channel sequences for potential nuclear localization signals using the program cNLS mapper [53]. The King (KJ756351) [27] and Bellono (KY355737) [29] sequences both contain a bipartite NLS, with different intervening linkers. These are KRIKKCGCKRLQDENPSVLSPKKKQRNG for King and KRIKKCGCKRPRYGYNGYLSTIQDENPSVLSPKKKQRNG for Bellono (note that Bellono has a 12 amino acid insert that is unique to their sequence: PRYGYNGYLSTI. These closely related, alternatively spliced bipartite NLS sequences are present at splice site 4 [38]. A 59-amino acid STREX exon alternatively spliced into this location [54] would substantially increase separation between the two clusters of basic K potentially attenuating functionality of the NLS. We analyzed BK channel proteomics and noted that the SVLSP sequence, or STLSP in human (NP_001309759.1), is frequently phosphorylated [55].

We used BlastP to search across species for proteins with sequence homology to the skate *kcnma1* bipartite NLS (Supplemental Table). We found that the bipartite NLS with variable intervening linker in *kcnma1* is highly conserved across diverse species. Vertebrates contain two clusters of basic amino acids, KRIKKCGCKR* upstream (* marks alternative splice 4, also known as c2) and KKKQRNG downstream. These sequences are absent in invertebrates, although Drosophila exhibits some homologous basic amino acids in these positions [34].

We evaluated this bipartite NLS for its ability to confer nuclear localization to skate BK channels transiently expressed in HEK293 cells. Skate *kcnma1* cDNA cloned into pcDNA3.1 mammalian expression vector was a gift from Nicholas Bellono and David Julius [29]. Site directed mutations were generated within the bipartite NLS, by InFusion cloning. In the proximal portion of the bipartite NLS, two basic amino acids were mutated to alanine, KR→AA. Separately, two serine residues within the linker domain that are sites of phosphorylation were mutated to alanine, SVLS→AVLA. HEK293 cells were either nontransfected, or transiently transfected with *kcnma1* wild type (WT) expression vector, KR→AA or SVLS→AVLA mutations. After 40 h cells were fixed with paraformaldehyde, permeabilized with TX-100 and immunostained for BK channels (Slo1 antibody) and subcellular markers, Lamin B1 or Calnexin. Confocal microscopy was performed to characterize the subcellular localization of transfected, intracellular BK channels

Compared to control cells, HEK293 cells transfected with BK channels exhibited increased BK immunoreactivity in plasma membranes and intracellular compartments (Figure 5, 6). Approximately 5-10% of all transfected cells showed high levels of intracellular BK expression. HEK293 cells expressing WT BK channels showed substantial colocalization with DAPI and lamin B1, consistent with nuclear translocation (Figure 5F-J). In contrast, HEK293 cells transfected with KR→AA or SVLS->AVLA mutations showed high expression of BK channels adjacent to but not overlapping nuclei stained with DAPI and lamin B1 (Figure 5K-O and P-T). We note that accumulation of peri-nuclear-excluded BK channels caused indentations of the nuclei (arrowheads in Figure 5L,Q; Figure 6 L,Q).

**Figure 5.**
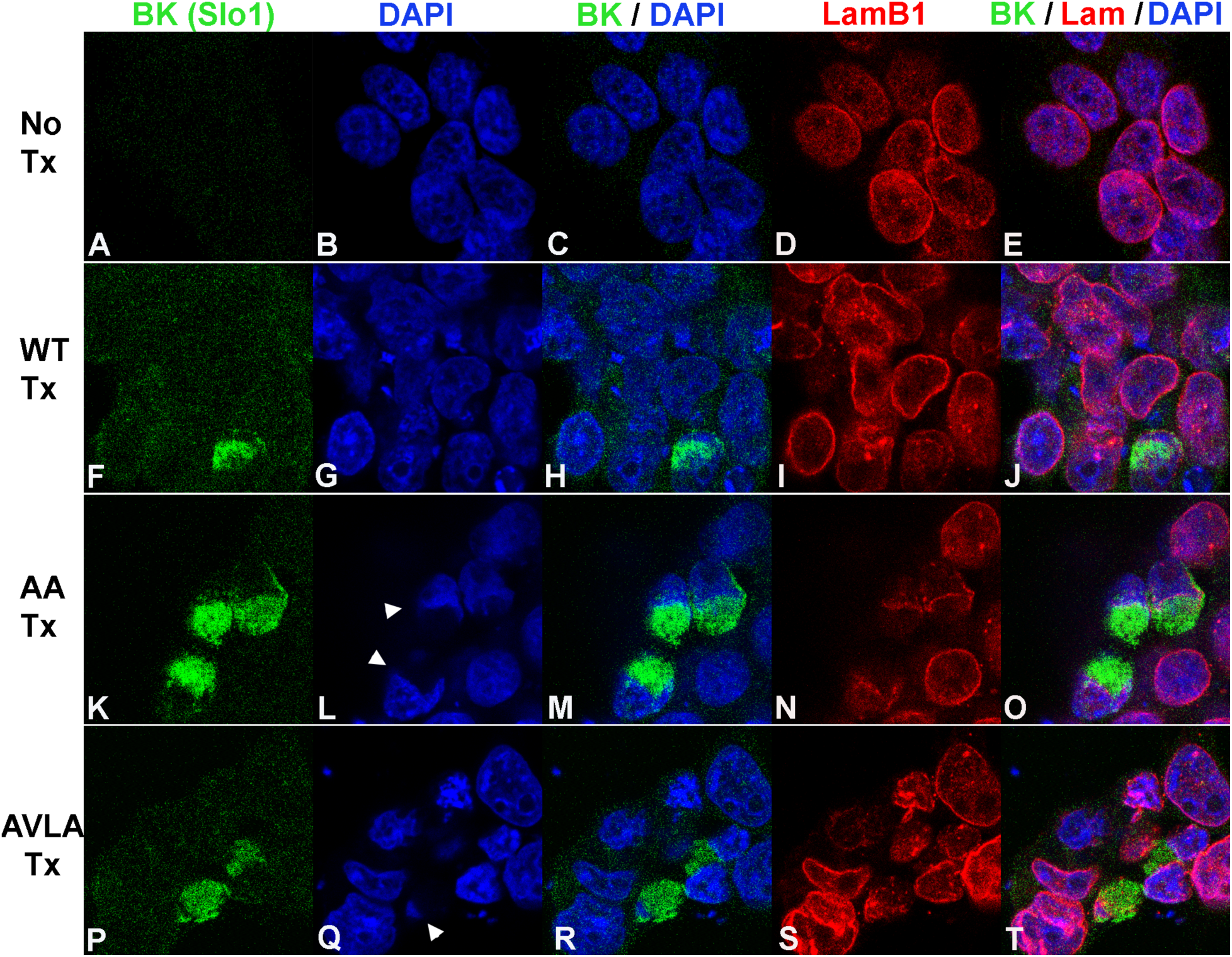
Localization of transfected skate BK channels and Lamin B1 in HEK293 cells. A-E: No transfection. A) anti-BK (Slo1 mouse monoclonal). B) DAPI. C) Merge of A and B. D) Lamin B1. E) Merge of A, B and D. F-J: WT transfection. F) anti-BK (Slo1). G) DAPI. H) Merge of F and G. I) Lamin B1 (rabbit polyclonal). J) Merge of F, G and I. K-O: AA mutation transfection. K) anti-BK (Slo1). L) DAPI. M) Merge of K and L. N) Lamin B1. O) Merge of K, L and N. P-T: AVLA mutation transfection. P) anti-BK (Slo1). Q) DAPI. R) Merge of P and Q. S) Lamin B1. T) Merge of P, Q and S.

**Figure 6.**
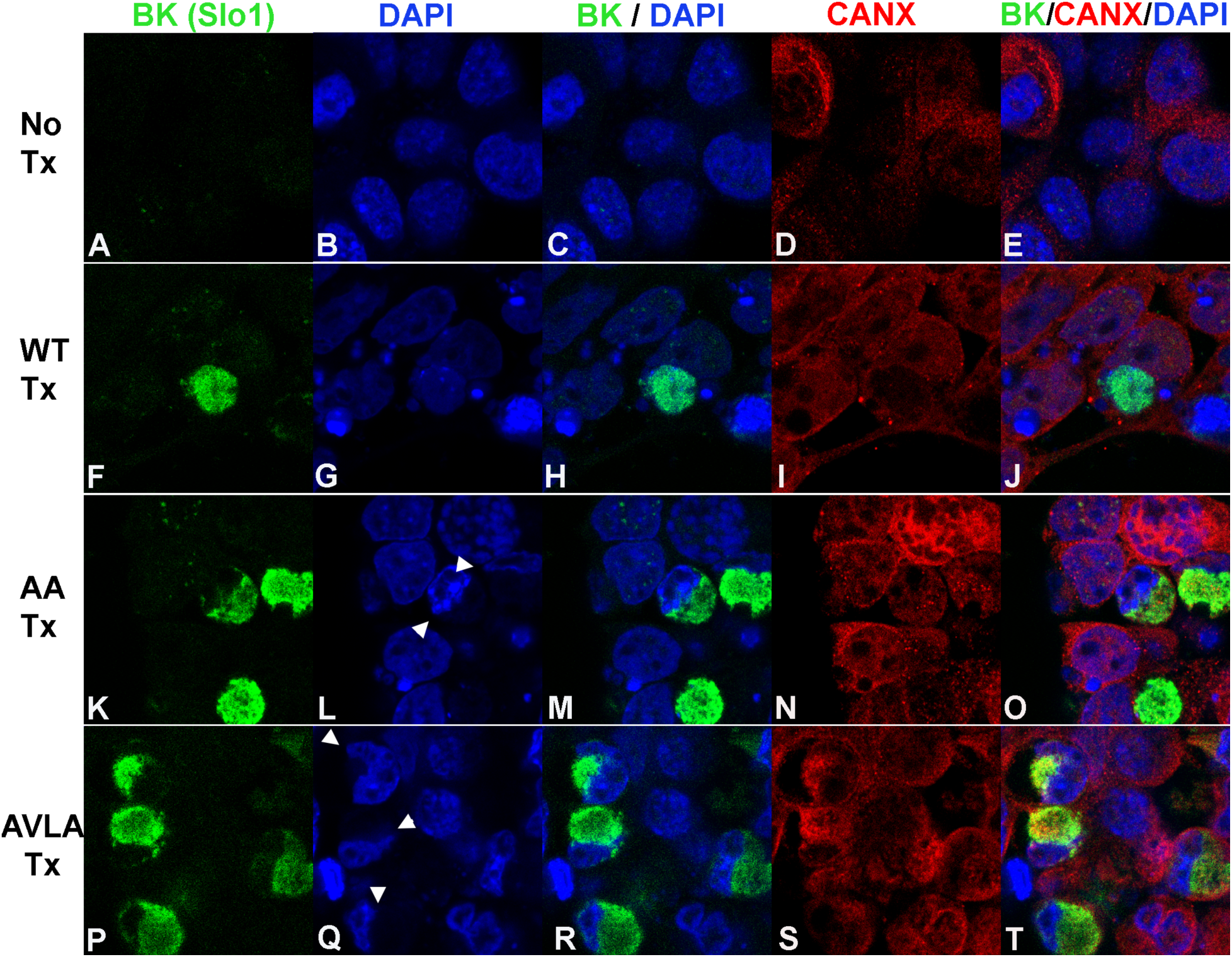
Localization of transfected skate BK channels and Calnexin in HEK293 cells. A-E: No transfection. A) anti-BK*α* (Slo1). B) DAPI. C) Merge of A and B. D) Calnexin. E) Merge of A, B and D. F-J: WT transfection. F) anti-BK (Slo1, mouse). G) DAPI. H) Merge of F and G. I) Calnexin (rabbit polyclonal). J) Merge of F, G and I. K-O: AA mutation transfection. K) anti-BK (Slo1). L) DAPI. M) Merge of K and L. N) Calnexin. O) Merge of K, L and N. P-T: AVLA mutation transfection. P) anti-BK (Slo1). Q) DAPI, R) Merge of P and Q. S) Calnexin. T) Merge of P, Q and S.

High intracellular expression of BK channels adjacent to nuclei suggested localization in endoplasmic reticulum (ER). Calnexin is an integral membrane protein chaperone that localizes to and is used as a marker of the ER [56]. BK-transfected HEK cells were immunostained with anti-BK and anti-Calnexin antibodies (Figure 6). HEK293 cells expressing WT BK channels showed higher levels of colocalization with DAPI than with calnexin, consistent with nuclear translocation (Figure 6 F-J). In contrast, HEK293 cells transfected with KR→AA or SVLS->AVLA mutations showed substantial colocalization with calnexin, and not with DAPI, consistent with retention of BK channels in the ER (Figure 6, K-O and P-T). Again, we noted that accumulation of peri-nuclear-excluded BK channels caused indentations of the nuclei (Figure 6Q, S).

We counted HEK293 cells with high intracellular expression of BK channels and determined the subcellular localization pattern in relationship to WT, KR→AA or SVLS->AVLA mutations. The results are shown in Figure 7. WT BK channels translocate into the nucleus at 69% frequency, whereas mutations KR→AA or SVLS→AVLA significantly reduced nuclear translocation to ∼15%. Compared to WT-transfected cells, SVLS→AVLA mutant transfected cells showed higher expression of intracellular BK channels (∼10%), and greater survival after transient transfections.

**Figure 7.**
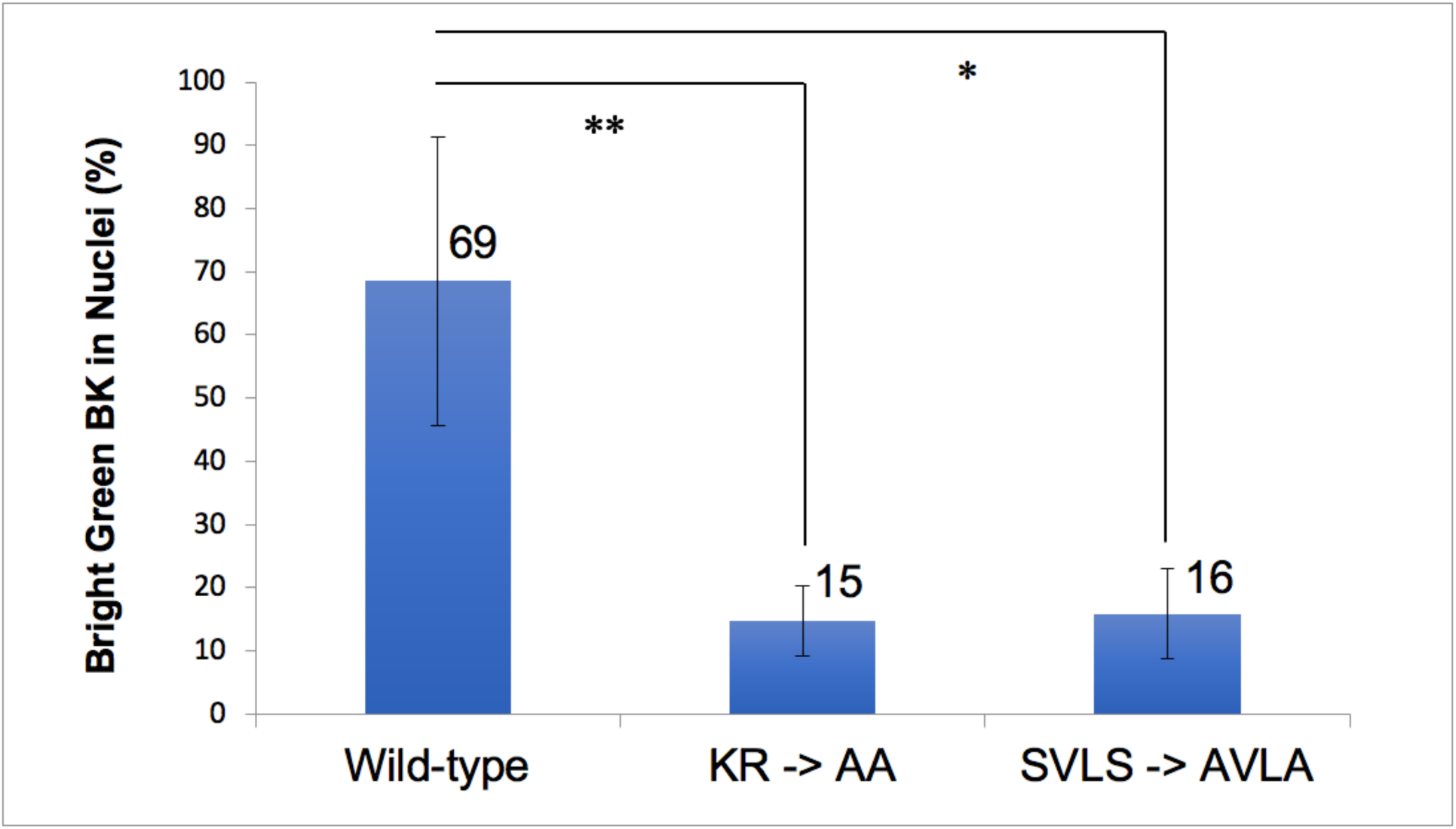
Localization of transfected skate BK channels in HEK293 cells. Significance of the differences between Wild Type, AA or AVLA mutation transfections was determined using one-way ANOVA with Tukey correction. p < 0.05 (*), < 0.01 (**).

We conclude that skate *kcnma1* BK*α* channel contains a bipartite NLS, in which specific mutations attenuate nuclear translation.

## Discussion

Confocal microscopy of skate ampullae unexpectedly revealed strong expression of BK channels in nuclei of electroreceptor cells. Across an alveolus we detected different patterns of BK channel expression: cells with strong nuclear staining showed light plasma membrane expression and reciprocally, cells with prominent plasma membrane expression showed low levels of nuclear expression. We infer that BK channels may partition between plasma membranes and intracellular membranes such as nuclear envelopes, nucleoplasmic reticulum [52] or endoplasmic reticulum. In elegant studies, Kv10.1 channels were shown to be inserted into plasma membranes and available for surface biotinylation by extracellular BirA enzyme and subsequently trafficked to inner nuclear membranes where they were determined by single channel recording to be functional [33]. We suggest by analogy that electroreceptor BK channels located in intracellular membrane compartments and plasma membranes are likely to be electrically functional.

Examination of the skate *kcnma1* sequences using cNLS mapper software identified clusters of basic amino acids likely to operate as a bipartite or two monopartite NLS. Notably the bipartite NLS occurs across splice site 4 [38], also identified as c2 [36]. Insertion here of alternative exons including the STREX exon, increases the separation between the two clusters of basic amino acids, and may disrupt the function as an NLS. A cross-species comparison of *kcnma1* sequences demonstrates high conservation of the bipartite NLS in all vertebrates. Additionally, there is high species conservation of the phosphorylation substrates threonine and serine immediately before the downstream basic portion of the bipartite NLS: <u>ST</u>L<u>S</u>PKKKQRNG. Membrane embedded proteins may be transported to the inner nuclear membrane through interactions of internal NLS with karophyrins (reviewed by [57]). The “SRKR” exon insertion at splice site 1, modulated by circadian rhythm, affects plasma membrane BK channel activation [37]. This insertion adds three basic amino acids 84 residues upstream of the mouse *kcnma1* bipartite NLS and creates an additional NLS that might affect nuclear trafficking; however, BK intracellular localization was not characterized.

Heterologous expression of *kcnma1* BK*α* subunits in HEK293 cells is an established method for characterizing electrophysiologic properties of BK*α* splice variants and functional consequences of additional BK beta subunits. Here, we used heterologous expression of skate *kcnma1* wild type [29] or mutated cDNAs in HEK293 cells to characterize intracellular BK*α* channel localization and regulation by the putative NLS. BK channel trafficking is undoubtedly complex and likely to depend on multiple addresses and post-translational modifications. For this study we chose not to fuse fluorescent proteins to BK*α* to minimize potential misfolding or mis-trafficking of the chimeric proteins. We performed BK localization analysis in *kcnma1*-transfected HEK293 cells after fixation, permeabilization and immunostaining similar to the skate ampulla. Our results establish that a fraction of transfected skate BK channels are expressed in the nucleus of HEK293 cells. Mutation of either, two basic amino acids in the first portion of the NLS or two serine residues upstream of the second portion of the NLS, independently attenuated nuclear localization of BK channels.

Numerous questions are raised by this study. What functional role may be served by BK channels in nuclei? Do nuclear BK channels contribute to calcium-regulated gene expression, if yes, which genes are most directly affected? Li et al [51] showed that BK channels in the nuclear envelope of hippocampal cells are functional and sensitive to pharmacologic inhibition by paxilline. Paxilline treatment of isolated nuclei caused depolarization of the nucleoplasm relative to the perinuclear lumen. Nuclear BK channels regulated the influx of calcium from the perinuclear space into the nucleoplasm. Paxilline-inhibition of nuclear BK channels produced transient increases in calcium in the nucleoplasm, and this nuclear calcium signaling affected calcium-dependent gene transcription, neuronal activity and dendritic arborization.

What regulates the partitioning of BK channels between nuclear envelopes and plasma membranes? Hutchinson-Gilford progeria syndrome (HGPS) manifests as premature aging and is caused by genetic mutations in the nuclear envelope protein lamin A. Electrophysiologic studies of human dermal fibroblasts from HGPS patients exhibited higher plasma membrane expression and larger outward potassium currents than from healthy young subjects [58]. These results suggest that functional BK channels that normally partition between nuclear envelopes and plasma membranes are forced into plasma membranes when nuclear envelopes are genetically disrupted. Dysregulation of plasma membrane BK channel expression may affect cell electrical excitability, influencing signal pathways important in aging, growth and proliferation [59].

How does plasma membrane expression of BK channels affect cell excitability and electroreceptor oscillation frequency and tuning? Miranda-Rottman et al demonstrated positional gradients of BK alternative splicing in auditory hair cells along the chicken cochlea. Our study suggests that alternative splicing at site 4 within the bipartite NLS may regulate the balance between BK channel nuclear localization and plasma membrane density thereby influencing cell excitability and frequency responsiveness [42].

How dynamic is the trafficking, and recycling of specific BK channels between nuclei and plasma membranes? Kinocilia are specialized structures scaffolded by microtubules and enveloped by plasma membrane that contain extremely high densities of signaling receptors including ion channels [60, 61]. The geometry of the cilium leads to a volume 5000-fold lower than the cytoplasm and enables signal amplification and quantal detection in sensory transduction. Cilia are anchored to basal bodies which operate as microtubule organizing centers, similar to centrioles [62]. Joukov et al [63] describe a physical connection between basal body apparatus and nucleus-associated microtubule organizing center as a means for communication between extracellular and intracellular domains. Our study suggests that BK channel partitioning between plasma membranes, including cilia, and nuclear envelopes may be regulated dynamically through posttranslational modifications within the bipartite NLS.

If BK trafficking is regulated by phosphorylation/dephosphorylation at the SVLS sequence in the bipartite NLS, what enzymes may be involved in these posttranslational modifications? Phosphorylation of Ser385 in the EBNA-1 NLS promoted nuclear import by increasing its affinity for karophyrin importin *α*5 [64]. What activates these enzymes? Are these enzymes potentially novel drug targets that might modulate BK trafficking and thereby cell signaling?

## Materials and Methods

### Dissection and whole mount immunostaining

Adult skates (*Leucoraja erinacea*) were purchased from Marine Biological Labs in Woods Hole, MA and shipped alive in chilled seawater. Animals were killed instantly by pithing of the brain and spinal cord. The ampullae were dissected with scissors from behind the gill slits. Approximately 200 ampullae with a short section of ampullary canal and afferent nerve were dissected in PBS under a stereomicroscope and cleaned of connective tissue. Ampulla were fixed in 1% paraformaldehyde in PBS for 1 h, then rinsed two times in PBS and stored for up to 1 week.

Whole mount staining was performed in 0.5 ml microfuge tubes in 100 microliters of solution, then blocked was with 10% FBS and 1% Triton X-100 detergent. Staining was performed with mouse and rabbit primary antibodies at final concentrations of 4 to 20 µg/ml in 0.1% FBS and 0.1% TX-100 for 48 h, followed by 3 serial washes with PBS −0.1% TX-100. Secondary antibodies were applied at 1:100 dilution; anti-mouse IgG(H+L) or IgG2*α* Alexa 488 or anti-rabbit Alexa 594. Nuclei were counterstained with DAPI.

Primary antibodies used were anti-BK*α*: KCNMA1 PA1-923 Invitrogen /ThermoFisher, rabbit polyclonal IgG, raised against synthetic peptide of human KCNMA1: T_945_ ELVNDTNVQFLDQDDD_961,_ skate epitope is identical at 17/18 positions, underlined: <u>TELVND</u>S<u>NVQFLDQDDD</u>. MaxiK*α* (B-1) sc-374142 Santa Cruz Biotechnology, mouse monoclonal IgG2b immunogen amino acids 937-1236 at C-terminus of human MaxiK*α*. Anti-Slo1, clone L6/60 MABN70 Millipore, mouse monoclonal IgG2a, immunogen is recombinant protein corresponding to segment S9-S10 of mouse Slo1 (NP_034740). For Ca_V_1.3 *cacna1d:* LS-B4915 LifeSpan Biosciences, mouse monoclonal IgG2a, immunogen is fusion protein encoding amino acids 8590875 of rat Ca_V_1.3 (Accession P27732): DNK------VTIDDYQEEAEDKD, skate epitope is conserved at 11 of 16 positions, underlined: <u>D</u>R<u>K</u>ILTGTQ<u>V</u>S<u>IDD</u>-<u>Q</u>--D<u>EDKD</u>. Lamin A/C: NBP2-59937 Novus Biologicals, mouse monoclonal IgG3. Lamin B1 Ab16048 AbCam, immunogen synthetic peptide corresponding to mouse Lamin B1 amino acids 400-500 conjugated to keyhole limpet hemocyanin. Calnexin HPA009433 Sigma Atlas Prestige Antibodies, rabbit polyclonal IgG, https://www.proteinatlas.org/ENSG00000127022-CANX/antibody.

Confocal imaging was performed using Zeiss LSM 880. Objective lenses used were apochromatic 10x/NA 0.45 and 40x/NA 1.2 glycerol immersion infinity/0.15-0.19. Z-stacks of up to 100 microns were collected, using separate tracks to avoid cross contamination of colors. Image files were processed using Fiji (Image J, NIH).

### Site-directed mutagenesis

Skate BK channel in mammalian expression vector pcDNA 3.1 – *kcnma1* was a generous gift from Bellono and Julius [29](Accession KY355737.1). Site directed mutations were generated using InFusion cloning system (ThermoFisher) with designated primers (nucleotide changes from WT in capital letters): KCNMA1_NLSmuKR655AA_1954plus 5’-tgtgggtgtGCAGCAccaagatatggctataatggatatctcagcac, and KCNMA1_NLSmuKR655AA_1962minus 5’-GGTGCTGCacacccacatttcttaattctctttgtatctgtg; KCNMA1_NLS_SVLS674AVLA_2017plus 5’-ccaGcagtgctgGctcccaaaaaaaagcaacggaacgggggc, and KCNMA1_NLS_SVLS674AVLA_2033minus 5’-ggagCcagcactgCtgggttttcatcttgaattgtgctgag. Mutagenesis was confirmed by DNA sequencing. WT; KR→AA mutation and SVLS→AVLA mutations.

### Immunofluorescence staining

HEK293 cells were transfected on multiwell imaging slides. Between 300-700 ng of DNA plasmid was mixed with JetPrime lipid transfection agent and applied to cells at 50% confluency for ∼40 h. Cells were fixed with 1% PFA, and immunostained at 10 µg/ml using anti-BK mouse monoclonal anti-Slo1 and rabbit polyclonal anti-Lamin B1 (nuclear envelope marker) or anti-Calnexin (ER marker [56]) antibodies. Non-transfected cells did not demonstrate prominent Slo1 staining. Transfected cells with prominent (intracellular) Slo1 staining were counted, and the fraction of those cells that exhibited overlap with DAPI nuclear stain was counted manually. The effect of site-directed mutations on nuclear localization of skate BK channels was determined quantitatively.

## Acknowledgements

We acknowledge the contributions of Alexis Froistad acquiring the initial confocal images of the skate ampulla.

